# Tissue Forge: Interactive Biological and Biophysics Simulation Environment

**DOI:** 10.1101/2022.11.28.518300

**Authors:** T.J. Sego, James P. Sluka, Herbert M. Sauro, James A. Glazier

## Abstract

Tissue Forge is an open-source interactive environment for particle-based physics, chemistry and biology modeling and simulation. Tissue Forge allows users to create, simulate and explore models and virtual experiments based on soft condensed matter physics at multiple scales, from the molecular to the multicellular, using a simple, consistent interface. While Tissue Forge is designed to simplify solving problems in complex subcellular, cellular and tissue biophysics, it supports applications ranging from classic molecular dynamics to agent-based multicellular systems with dynamic populations. Tissue Forge users can build and interact with models and simulations in real-time and change simulation details during execution, or execute simulations off-screen and/or remotely in high-performance computing environments. Tissue Forge provides a growing library of built-in model components along with support for user-specified models during the development and application of custom, agent-based models. Tissue Forge includes an extensive Python API for model and simulation specification via Python scripts, an IPython console and a Jupyter Notebook, as well as C and C++ APIs for integrated applications with other software tools. Tissue Forge supports installations on 64-bit Windows, Linux and MacOS systems and is available for local installation via conda.

**Author Summary:** Tissue Forge is a physics-based modeling and simulation software environment for research problems in physics, chemistry and biology. Tissue Forge supports modeling at a wide range of scales, from as small as the sub-nanometer, to as large as hundreds of micrometers, using particle-based models. It provides rich features for simulation development and application at all stages of model-based research, like real-time simulation visualization and interactivity, and off-screen batch execution, rendering, and GPU acceleration. Users can employ built-in models to represent a wide variety of physical processes, like chemical reactions, fluid convection and intercellular adhesion, or define their own models for agentand rule-based modeling. Tissue Forge is open-source, free and easy to install, supports simulation development in C, C++ and Python programming languages, and can be used as integrated software or in an interactive IPython console and Jupyter Notebook. Tissue Forge also provides a dedicated space for application-specific and user-contributed modeling and simulation features, and developers are welcome to contribute their custom features for distribution in future releases.

## 2 Introduction

Computational modeling and simulation are key components of modern biological research. Simulations codify knowledge into computable representations that can challenge and validate our understanding of complex biological processes. A well defined model not only explains currently available data but also predicts the outcomes of future experiments. Biological computer simulations can address a wide range of length scales and employ numerous numerical and simulation technologies. Scales include that of the atomic bond length to model small molecules, proteins and other biological macromolecules, the macromolecular scale to model protein aggregates, the subcellular and cellular scales to model cells and aggregates of cells, the tissue scale to model long-range interaction between cell aggregates that give rise to organ-level behaviors, the whole-body scale where organs interact, and the population scale where individuals interact with each other and their environment. At various biological scales, models can represent biological objects as either discrete or as numerically aggregated populations, and so different mathematical and computational approaches are used to simulate behaviors at each scale. When spatiality is explicitly modeled, molecular dynamics (**MD**) simulations are often used at the atomic and macromolecular scales and spatial agent-based models are used at the higher scales. Often, discrete biological objects (*molecules, cells, cell aggregates*) are appropriately modeled as discrete objects at a particular scale, and then as numerically aggregated populations at higher scales using continuous dynamics like ordinary differential equations (**ODEs**) and partial differential equations (**PDEs**), which then describe the dynamics of a population of objects. For example, modeling at the multicellular scale can represent molecules of a given chemical species as densities or amounts, and at the molecular level as discrete molecules. While population models can have significant explanatory value, biology is intrinsically spatial. Emergent biological properties and behaviors arise in part because of the spatial relationships of their components. Population models sacrifice this aspect of biological organization.

In the subcellular, cellular and multicellular modeling domain, most spatiotemporal agent-based biological simulation tools only support one cellular dynamics simulation methodology, and focus on a particular problem domain with a particular length scale. For example, CompuCell3D (**CC3D**) [1] and Morpheus [2] implement cell model objects using the Cellular Potts model (**CPM**)/ Glazier-Graner-Hogeweg (**GGH**) formalism [3], and only support Eulerian, lattice-based models, while others like PhysiCell [4] and CHASTE [5] support modeling cells with Lagrangian, lattice-free, particle-based center models as simple, point-like cell particles. Lattice-free, particle-based methods can be extended to include subcellular detail using the Subcellular Element Model [6], which could support modeling the spatial complexity of cell shape, cytoskeleton and extracellular matrix. Extending the CPM/GGH to include cellular compartments [7] allows representation of subcellular components like the nucleus, critical molecular species or regions with specific properties but does not support specific representation of macromolecular machinery. Typically, modelers who are interested in subcellular and cellular detail must use and adapt general-purpose MD simulation tools like LAMMPS [8], HOOMD-blue [9], NAMD [10] or GROMACS [11]. For example, Shafiee et al., customized LAMMPS to model cells as clusters of particles to simulate spheroid fusion during spheroid-dependent bioprinting [12].

Most MD simulation tools are designed to parse and execute models that are theoretically well defined and MD simulation specifications and engines tend to be well optimized for computational performance. Most assume a fixed numbers of objects within a model and do not support runtime object creation, destruction or modification. Many do not support real-time simulation visualization and user interactivity. In addition, extending these modeling environments with custom modeling and simulation features requires software development in C or C++ code. Results can be post-processed after execution, though this requires developing a pipeline of model development, simulation execution and data generation using a simulation tool, and data visualization and analysis using different visualization tools (*e*.*g*., The Visualization Toolkit [13]) or a general purpose programming language like Python, which significantly increases user effort to produce useful results. To reduce user effort required to produce publishable simulation results and analysis, some simulation tools provide realtime simulation visualization and limited simulation interaction (*e*.*g*., CC3D and Morpheus). Cell simulation tools with real-time visualization are often implemented as stand-alone programs, rather than as portable libraries that support integration with other modeling environments. This lack of software interoperability also complicates using simulation tools with other specialized software libraries (*e*.*g*., optimization tools) in advanced computational workflows for solving difficult biological problems such as reverse-engineering model parameters, interrogation of mechanisms, or Bayesian modeling of populations.

This paper presents Tissue Forge, an open-source, real-time, modeling and simulation environment for interactive biological and biophysics modeling applications over a broad range of scales. Tissue Forge is designed to address many of the aforementioned issues and challenges. Tissue Forge enables agent-based, spatiotemporal computational modeling at scales from the molecular to the multicellular. It is designed for ease of use by modelers, research groups and collaborative scientific communities with expertise ranging from entryto advancedlevel programming proficiency. It supports all stages of model-supported research, from initial model development and validation to large-scale virtual experiments. Here we describe the philosophy, mathematical formalism and basic features of Tissue Forge. To demonstrate its usefulness across multiple disciplines in the physical and life sciences, we also present representative examples of advanced features at a variety of target scales.

## 3 Overview

Tissue Forge seeks address some of the limitations of current modeling packages by providing a spatiotemporal modeling and simulation environment that supports multiple lattice-free, particle-based methods for agent-based modeling. It simplifies research by supporting representation of a wide range of scales encountered in biophysics, chemistry and biological applications. Tissue Forge supports the development, testing and deployment of models in large-scale, highperformance simulation, performed by users with a wide range of expertise and coding proficiency in multiple programming languages.

### 3.1 Problem Domain

Simulation of complex systems, particularly in biological problems, is difficult for a number of reasons. Difficulties exist for both the domain knowledgeable modeler and the modeling tool developer. Problems in cell biology and biophysics applications often require representations of objects and processes at multiple scales, which resolve to spatiotemporal, agent-based models with complex rules and decision making using embedded models of internal agent state dynamics (*e*.*g*., chemical networks). Since such models are experimentally or empirically determined and highly diverse, their implementation requires flexible, robust model and simulation specification. Likewise, the spatial scale itself presents the challenge of choosing an appropriate mathematical framework for creating model objects and processes (*e*.*g*., whether to model a cell with complex shape or simply as a sphere). Often, the modeler must learn a new software tool for each spatial scale they wish to model. In addition, the model features and computational performance of a particular software tool can be limited by the underlying mathematical framework, unpermissive or demanding object definitions, or the need for efficient use of computing resources.

Tissue Forge addresses these issues by providing an agent-based, spatiotemporal modeling and simulation framework built on a flexible, particle-based formalism. Particles, which are the fundamental agents of any Tissue Forge simulation, are suitable basic objects in model construction because they minimally constrain a model description. A Tissue Forge particle is an instance of a categorical descriptor called a “particle type,” and is a discrete agent that has a unique identity, occupies a position at each moment in time and has velocity and mass or drag. Tissue Forge imposes no further restrictions on what physical or abstract object a particle represents. This framework has the theoretical and computational flexibility to enable agent-based, spatiotemporal computational models across a broad range of scales. An instance of a particle could represent an atom, or a cell, or a multicellular aggregate. Tissue Forge accommodates models with both preand user-defined particle behaviors and interactions, the creation and deletion of particles at runtime, and consistent object modeling at multiple scales.

#### Interactive and Batch Execution

Tissue Forge supports the efficient development agent-based models of complex systems. In general, the development of a computational model involving multiple interacting agents requires iterative cycles of model development, execution, analysis, and refinement. During model exploration, refinement and validation, modelers can benefit from a simulation environment that allow them to observe, interact with, and refine a simulation as it executes (*i*.*e*., real-time simulation and visualization). However, computationally intensive investigations of developed models (*e*.*g*., characterizing emergent mechanisms or the effects of system stochasticity, systems with large numbers of objects) require efficient high-performance computing utilization and batch execution. Tissue Forge supports both interactive and batch operation, providing both rapid and intuitive model development and high-performance simulation execution, so that modelers do not need to find and learn multiple software tools or settle for a tool that is either, but not both, feature rich or computationally efficient. Its interactive simulation mode is a stand-alone application with realtime visualization and user-specified events. Its batch mode leverages available resources in high-performance computing environments such as computing clusters, supercomputers, and cloud-based computing, and exports simulation data and high-resolution images. In batch mode, Tissue Forge can be included in workflows to carry out modeling task such as model fitting or simulation of replicates and populations.

#### Open Science Support

Development and dissemination of models that leverage interdisciplinary knowledge and previous modeling projects require robust support for scientific communication, collaboration, training and reuse. Tissue Forge provides a declarative model specification for many basic aspects of particle-based models and simulations (*e*.*g*., particle type definitions, particle interactions and stochastic motion via generalized force and potential definitions) with robust support for procedural specification of complex, agent-based models particular to specific applications. Tissue Forge also supports model sharing and collaborative development by providing built-in support for exporting and importing simulations and model object states to and from human-readable string data (using JSON format). In support of collaborative, community-driven and application-specific development of models, the Tissue Forge code base provides a designated space in which developers can implement features in customized Tissue Forge builds. Extending the Tissue Forge API with custom interfaces requires minimal effort in all supported software languages. Developers are also welcome to submit their custom features to the public Tissue Forge code repository for future public release as built-in features, or to design their software applications using Tissue Forge as an external software library. Along with executing scripted simulations specified in C, C++ and Python programming languages, Tissue Forge also supports collaboration, training and scientific communication through its Python API support for interactive simulations in Jupyter Notebooks. Tissue Forge simplifies robust model construction and simulation development through expressive model specification (*e*.*g*., process arithmetic), a flexible event system for implementing model-specific rules (*e*.*g*., agent rules) and simulation-specific runtime routines (*e*.*g*, importing and exporting data), and a simple, intuitive simulation control interface (*e*.*g*., switching between interactive and off-screen execution).

### 3.2 Concepts

Tissue Forge updates the trajectory of a particle in time by calculating the net force acting on the particle. Forces determine the trajectory of a particle according to the dynamics of the particle type. Tissue Forge currently supports Newtonian and Langevin (overdamped) dynamics, which can be individually specified for each particle type of a simulation.

For Newtonian dynamics, the position ***r***_*i*_ of the *i*th particle is updated according to its acceleration, which is proportional to its mass *m*_*i*_ and the total force ***f***_*i*_ exerted on it,

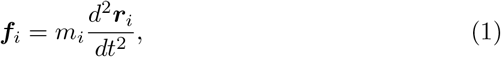

and for Langevin (overdamped) dynamics, *m*_*i*_ is the drag coefficient and the particle velocity is proportional to the total force,

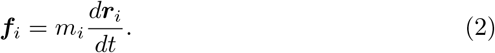

Tissue Forge supports three broad classes of force-generating interaction,

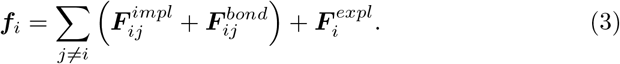

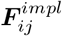 is the force due to *implicit* interactions between the *i*th and *j*th particles, 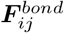 is the force due to *bonded* interactions between the *i*th and *j*th particles, and 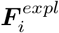 is the explicit force acting on the *i*th particle. Implicit interactions result automatically from interaction potentials between pairs of particles of given types. Bonded interactions act between specific pairs of individual particles (Figure 1A). Explicit forces act on particles through explicitly-defined force descriptions and do not necessarily represent inter-particle interactions (*e*.*g*., gravity, internal noise, system thermal equilibrium). Tissue Forge provides built-in forceand potential-based definitions, supports user-specified definitions for both, and permits applying an unlimited number of executable Tissue Forge force and potential objects to individual particles and particle types.

**Figure 1:**
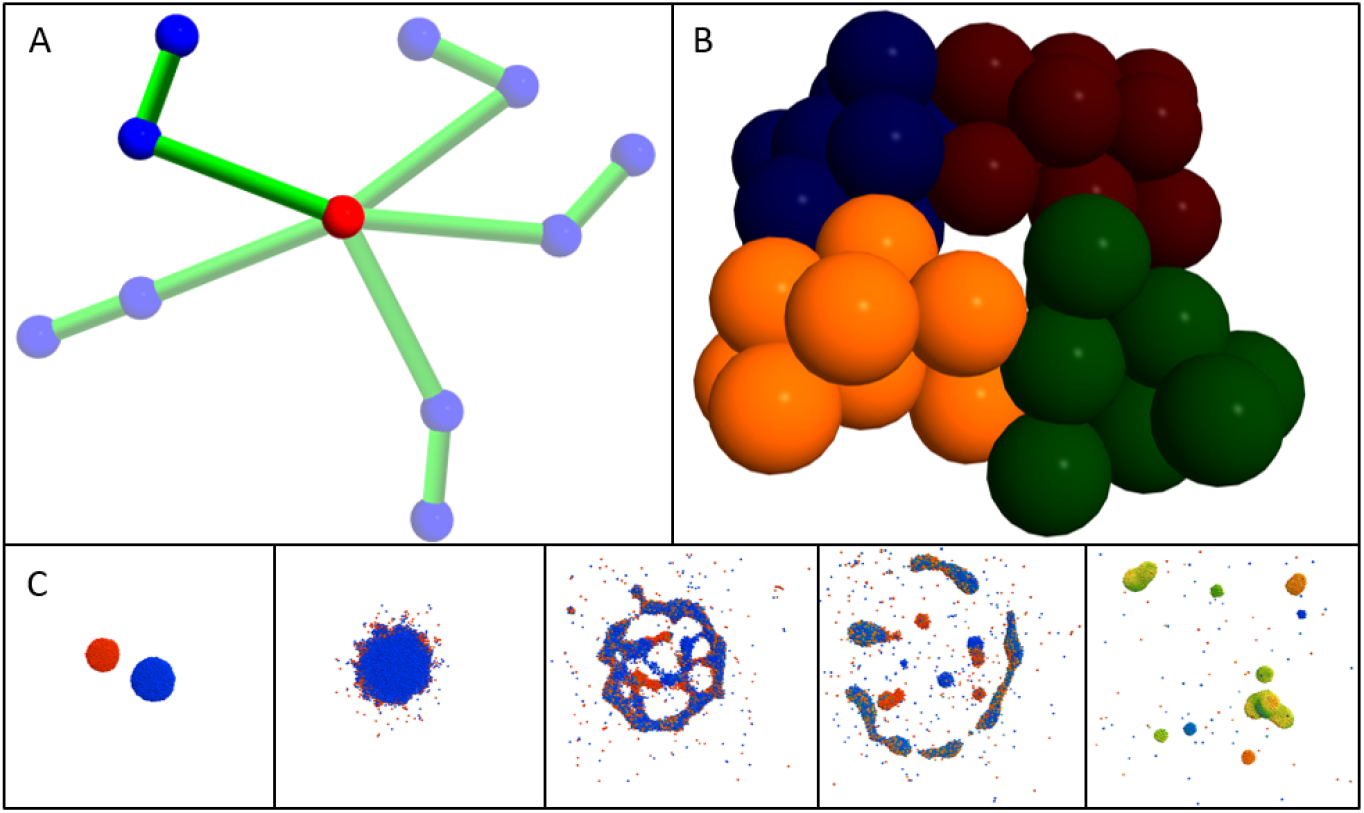
Examples of Tissue Forge modeling features. A: Five superimposed snapshots of a double pendulum implemented in Tissue Forge. Bonded interactions (represented as green cylinders) explicitly describe the interaction between a particular pair of particles, while a constant force acts on the blue particles in the downward direction. The red particle is fixed. B: Four Tissue Forge clusters representing biological cells, each consisting of ten particles whose color demonstrates cluster membership. Potentials describe particle interactions by whether they are in the same cluster (*i*.*e*., intracellular) or different clusters *i*.*e*., intercellular. C: Tissue Forge simulation of chemical flux during fluid droplet collision. Each particle represents a portion of fluid that carries an amount of a diffusive chemical, the amount of which varies from zero (blue) to one (red). When two droplets carrying different initial chemical amounts collide, resulting droplets tend towards homogeneous chemical distributions.

Implicit interactions are defined in Tissue Forge using potential functions and applied according to the types of two interacting particles. The force between the *i*th and *j*th interacting particles resulting from their implicit interactions is calculated as the sum of each *k*th potential 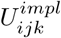 that defines the implicit interaction,

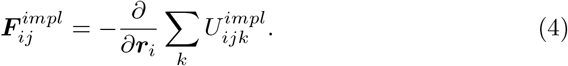

Bonded interactions are defined in Tissue Forge using potential functions and are applied according to the identities of two interacting particles. The force between the *i*th and *j*th interacting particles resulting from their bonded interactions is calculated as the sum of each *k*th potential 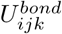 that defines the bonded interaction,

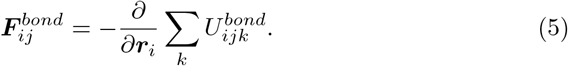

Explicit forces can be defined on the basis of particle type or on individual particles. The force on the *i*th particles resulting from external forces is calculated as the sum of each *k*th explicit force 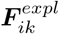

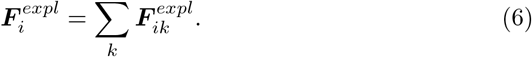

Since Tissue Forge enables the implementation and execution of models at different length scales, particles in a simulation may represents objects with a wide variety of possible behaviors. A particle could be atomic and subject to energy-conserving, implicit interactions (*e*.*g*., Coulomb, Morse or LennardJones potentials) as in classic MD. Particles can also represent portions of material that constitute larger objects (*e*.*g*., a portion of cytoplasm) and can carry quantities of materials within them (*e*.*g*., convection of a solute chemical in a portion of a fluid, Figure 1C). Tissue Forge provides built-in features to enable particle-based modeling and simulation of fluid flow based on transport dissipative particle dynamics (*tDPD*) and smooth particle hydrodynamics, including a predefined tDPD potential 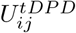 that can be applied when describing the interactions of a simulation,

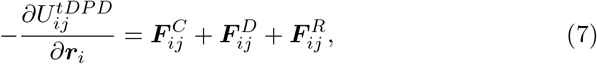

where the interaction between the *i*th and *j*th fluid-like particles is a sum of a conservative force 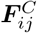, a dissipative force 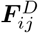 and a random force 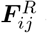 acting on the *i*th particle.

To support treating particles as constituents of larger objects, Tissue Forge provides a special type of particle, a *cluster*, whose elements can consist of constituent particles or other clusters. Clusters provide a convenient way to define implicit interactions that only occur between particles within the same cluster (*e*.*g*., intracellular interactions), called *bound* interactions, and those that only occur between particles from different clusters (*e*.*g*., intercellular interactions), called *unbound* interactions (Figure 1B).

To allow particles to carry embedded quantities, Tissue Forge supports attaching to each particle a vector of states that can evolve during a simulation. The values of the states can evolve according to laws defined between pairs of particle types for inter-particle transport (*e*.*g*., diffusion), which Tissue Forge automatically applies during simulation, or according to local, intra-particle reactions. The time evolution of a state vector ***C***_*i*_ attached to the *i*th particle is,

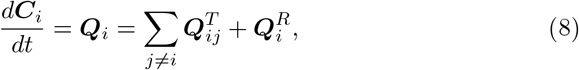

where the rate of change of the state vector attached to the *i*th particle is equal to the sum of the transport fluxes 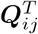 between the *i*th and each nearby *j*th particle and the local reactions 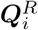.

### 3.3 Basic Features

Tissue Forge supports model and simulation specification using classes, objects and functions typical to object-oriented concepts in C, C++ and Python programming languages. In Python, custom Tissue Forge particle types can be defined by creating Python classes and specifying class attributes (Listing 1).

**Listing 1:**
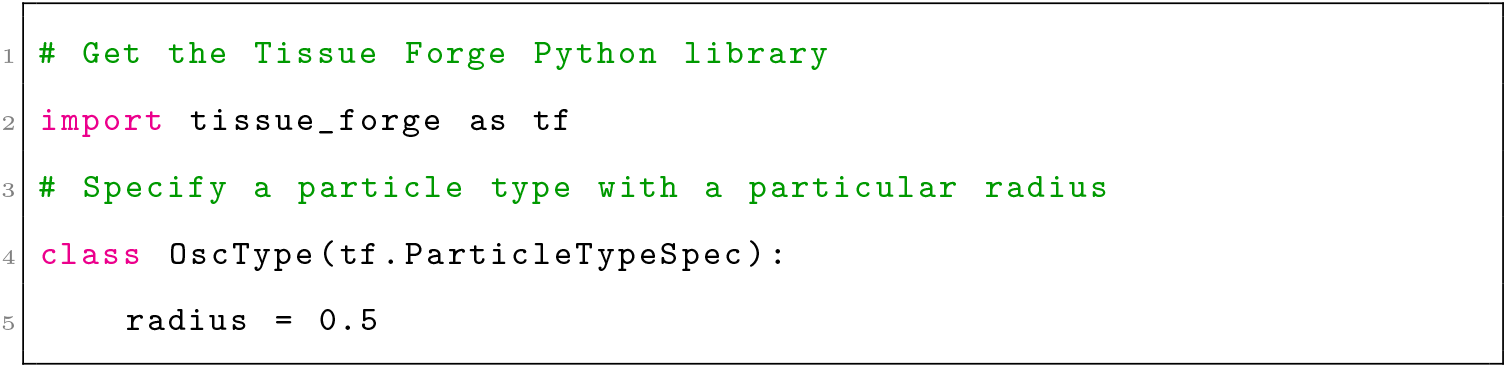
Importing the Tissue Forge library and declaring a particle type in Python. Comments are shown in green.

Tissue Forge allows specification of particle types without an initialized Tissue Forge runtime. However, initializing the Tissue Forge runtime, which in Python only requires a call to a single module-level function, permits retrieving template executable particle types that can be used to create particles (Listing 2). When a particle of a particular particle type is created, the particle inherits all attributes of its type (*e*.*g*., mass), which can in turn be modified for the particular particle at any time during simulation. Initializing the Tissue Forge runtime requires no user-specified information, in which case a default configuration is provided, but explicit initialization provides a number of customization options to tailor a simulation to a particular problem (*e*.*g*., domain size, interaction cutoff distance).

**Listing 2:**
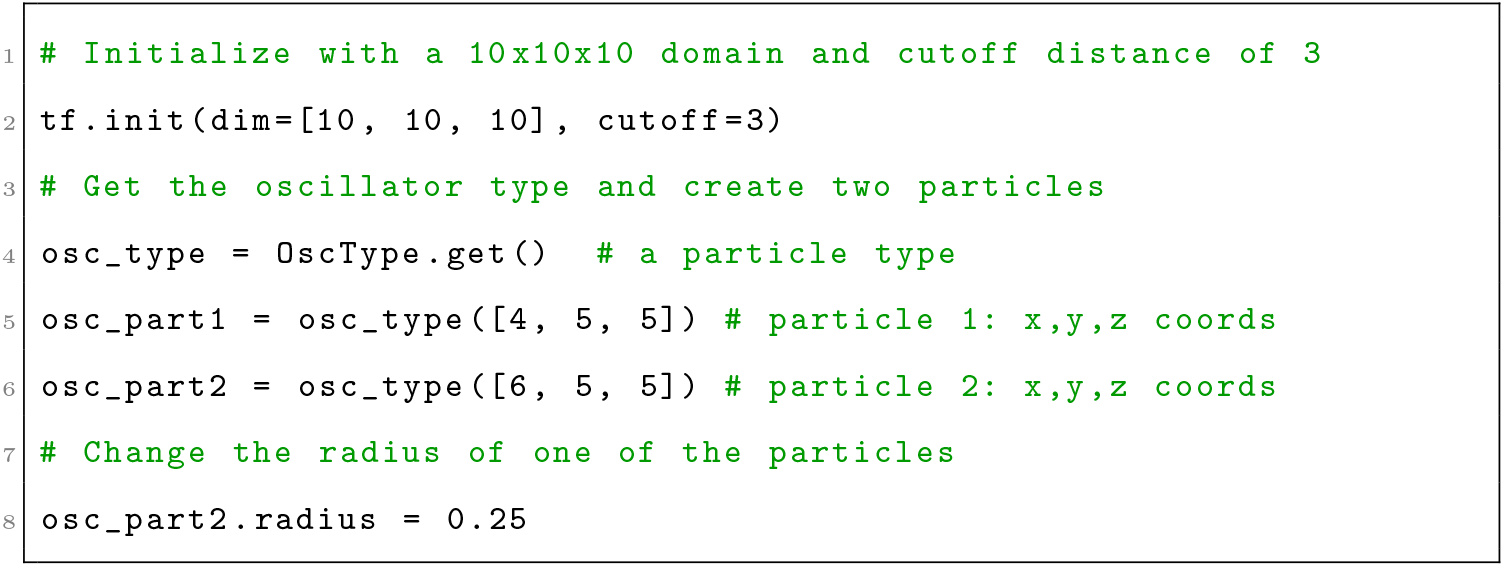
Initializing a Tissue Forge simulation, retrieving an executable particle type and creating particles in Python.

Users specify and apply interactions, whether using built-in or custom potential functions or explicit forces, by creating Tissue Forge objects that represent processes (*e*.*g*., a force object), called *process objects*, and applying them categorically by predefined ways that processes can act on objects (*e*.*g*., by type pairs for implicit interactions). Tissue Forge calls applying a process to model objects *binding*, which Tissue Forge applies automatically during simulation execution according to the model objects and process. For example, users can specify an implicit interaction between particles to two types by creating a potential object and binding it to the two particle types (Listing 3).

**Listing 3:**
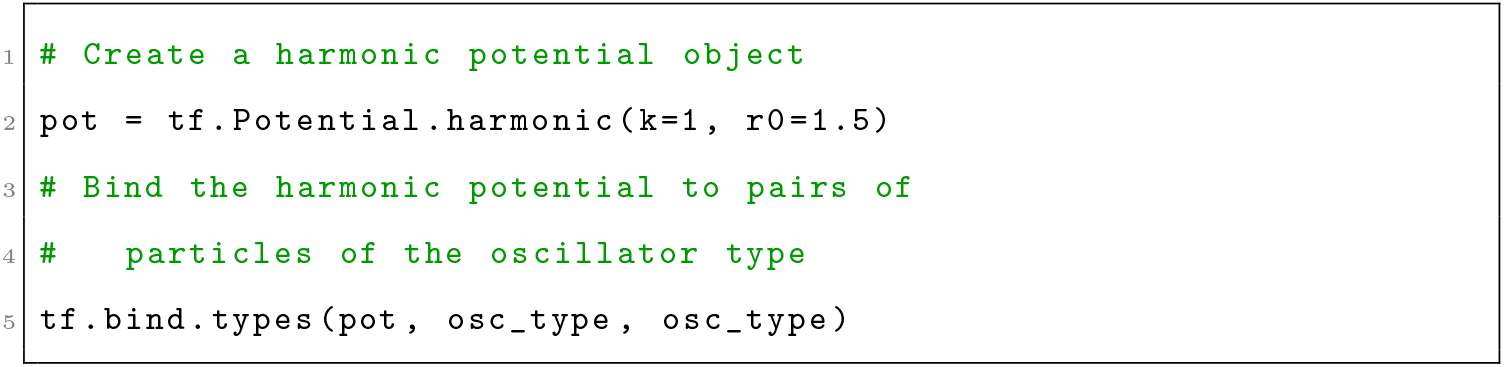
Creating a Tissue Forge potential and binding it to particles by type in Python.

Tissue Forge provides fine-grained simulation control, where each integration step can be explicitly executed, with other user-defined tasks accomplished between executing simulation steps (*e*.*g*., exporting simulation data). For interactive execution, Tissue Forge simulations are usually executed using a basic run function, which executes an event loop that (1) integrates the universe, (2) processes user input (*e*.*g*., keyboard commands), (3) updates simulation visualization, and (4) executes an event system with user-defined events. The Tissue Forge event system allows users to insert instructions into the event loop via user-defined functions (Listing 4). Events can be executed at arbitrary frequencies, can automatically retrieve simulation data (*e*.*g*., a randomly selected particle of a specific type), and can change qualities of individual particles (*e*.*g*., change the radius of a particular particle based on its environment).

**Listing 4:**
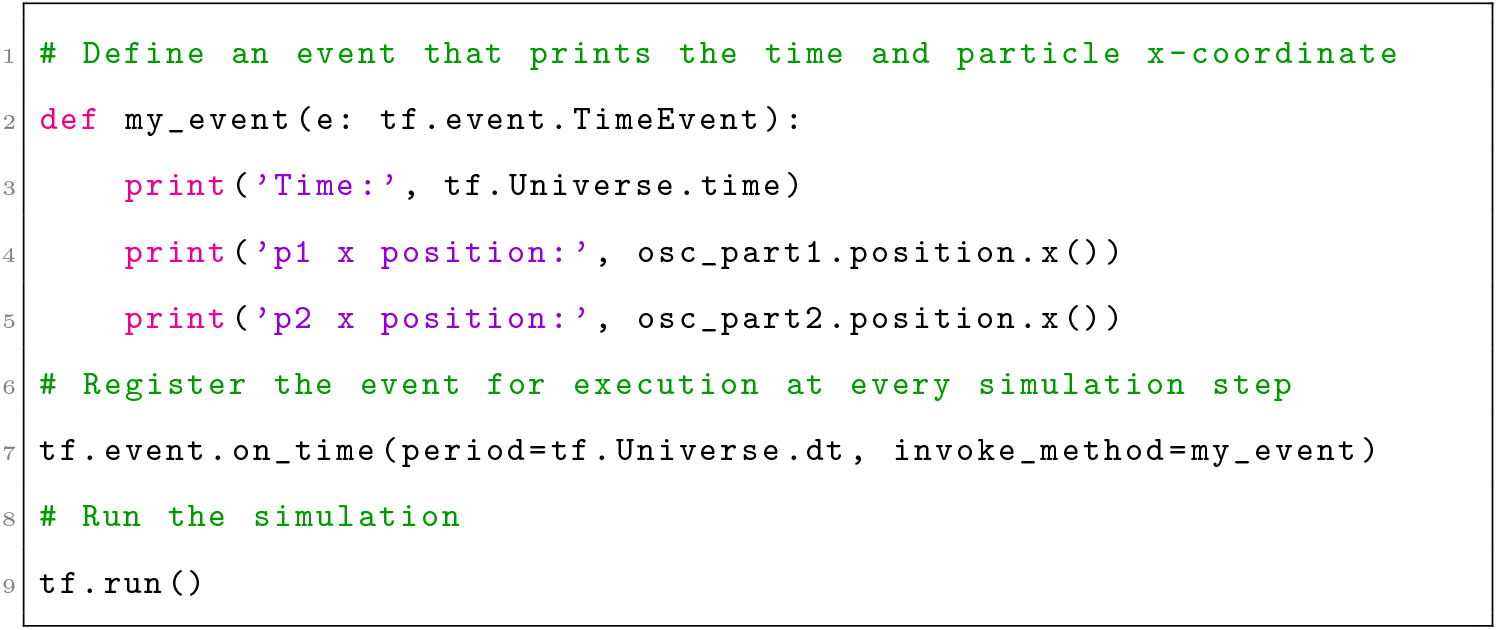
Creating a Tissue Forge event and running an interactive simulation in Python.

During simulation execution, including during execution of user-defined events, Tissue Forge objects are available for accessing and manipulating simulation, universe and system information. The Python code described in this section generates the Tissue Forge simulation depicted in Figure 2 (see Supplementary Materials S2), and also prints the current simulation time and *x*-coordinate of both particles at every simulation step. This simulation can be executed as a Python script or in an IPython console.

**Figure 2:**
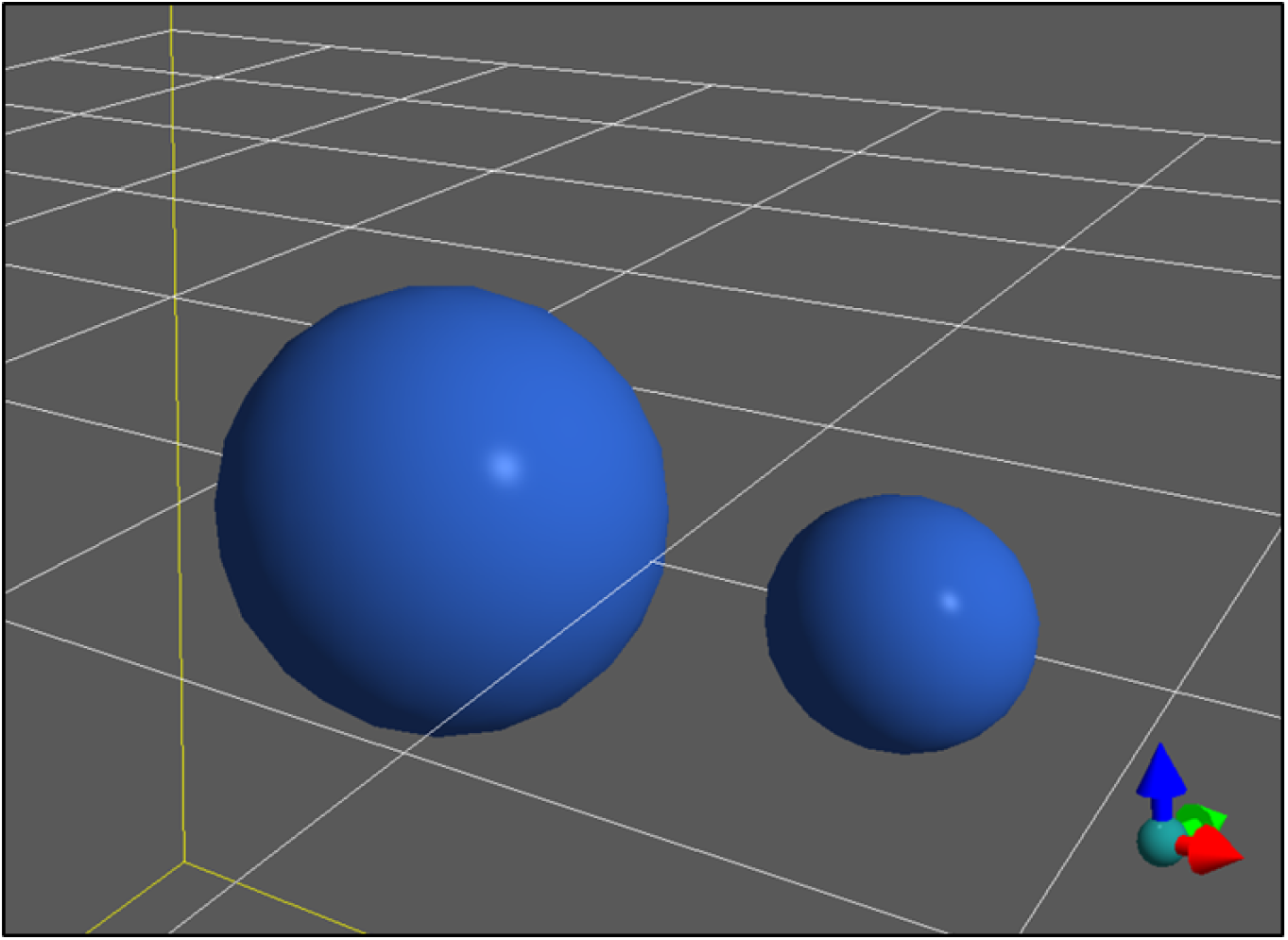
Tissue Forge simulation of a simple oscillator with two particles interacting via a harmonic potential. Tissue Forge helps to orient the user by drawing a yellow box around the simulation domain, a white grid along the *xy* plane at the center of the domain, and an orientation glyph at the bottom right to demonstrate the axes of the simulation domain with reference to the camera view, where red points in the *x* direction, green in the *y* direction and blue in the *z* direction.

**Figure 3:**
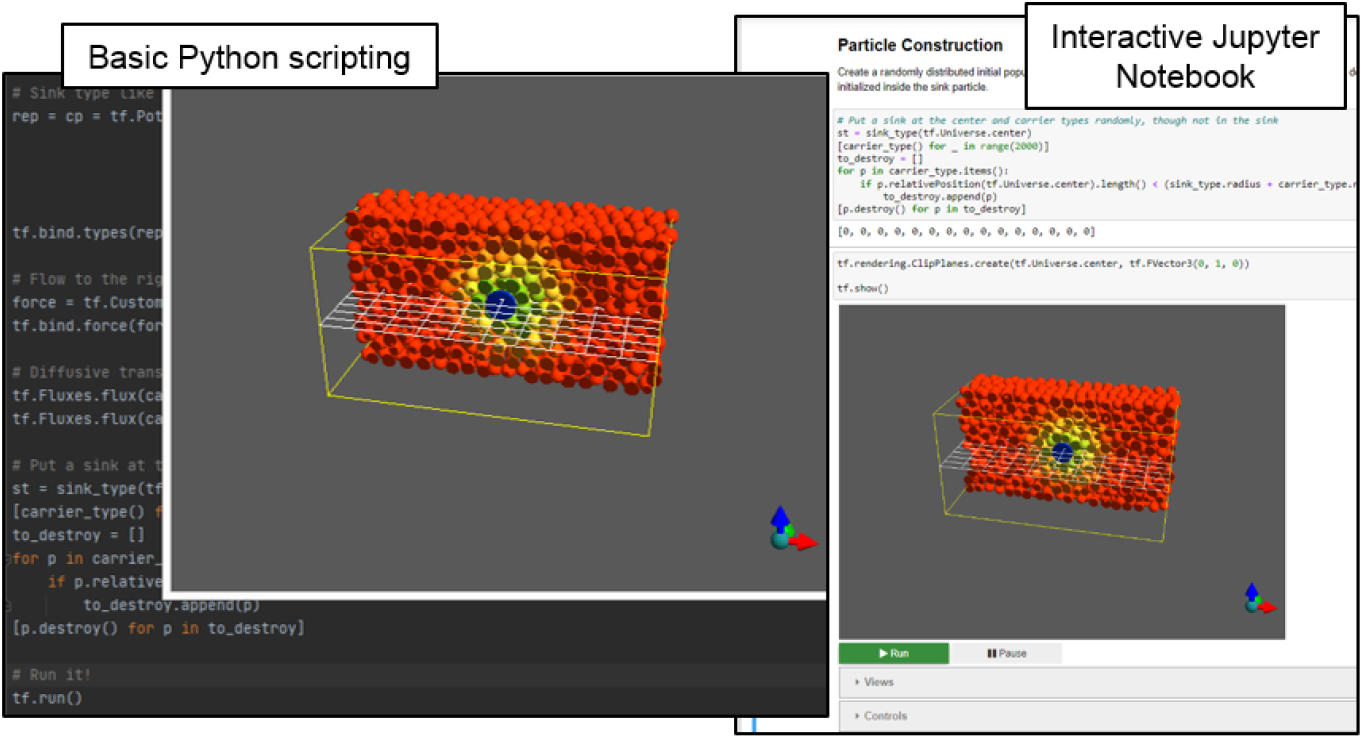
Sample use of the Python API to specify an interactive simulation of convection of a species near a species sink in a Python script (left) and in an interactive Jupyter Notebook (right).

In a Jupyter Notebook, this code executes the same simulation but generates an additional user interface, which provides widgets for interactive simulation controls, *e*.*g*., for pausing and resuming the simulation, and choosing predefined camera views (Figure 3). When running Tissue Forge from a Python script or IPython console, the interface supports mouse control (*e*.*g*., click and drag to rotate) and predefined and user-defined keyboard commands (*e*.*g*., space bar to pause or resume the simulation). In interactive contexts like IPython and Jupyter Notebooks, the Tissue Forge event loop recognizes user commands issued *ad hoc* during simulation, allowing on-the-fly modification of the simulation state, which is especially useful during model development and interrogation (*e*.*g*., when testing the effects of the timing of an event).

### 3.4 Sample Modeling Applications

Beyond the provided catalogue of built-in potentials, potential arithmetic (*e*.*g*., a potential object as the sum of two potential objects) and support for user-specified custom potentials, Tissue Forge provides process objects for binding potential-based process between specific particles (*i*.*e*., a *bonded* interaction). Bonded interactions are a key component of MD modeling. Tissue Forge provides a number of bond-like processes to apply potentials for various types of bonded interactions. Each bonded interaction has a representative object that contains information about the bonded interaction (*e*.*g*., which particles, what potential) that Tissue Forge uses to implement it during simulation. Currently, Tissue Forge provides the Bond for two-particle bonded interactions (where the potential is a function of the Euclidean distance between the particles, Figure 4, top left), the `Angle` for three-particle bonded interactions (where the potential is a function of the angle between the vector from the second to first particles and the vector from the second and third particles, Figure 4, top middle), and Dihedral (torsion angle) for four-particle bonded interactions (where the potential depends on the angle between the plane formed by the first, second and third particles and the plane formed by the second, third and fourth particles, Figure 4, top right). Like particles, all bonded interactions can be created and destroyed at any time during simulation, and bonded interactions can also be assigned a dissociation energy so that the bond is automatically destroyed when the potential energy of the bond exceeds its dissociation energy.

**Figure 4:**
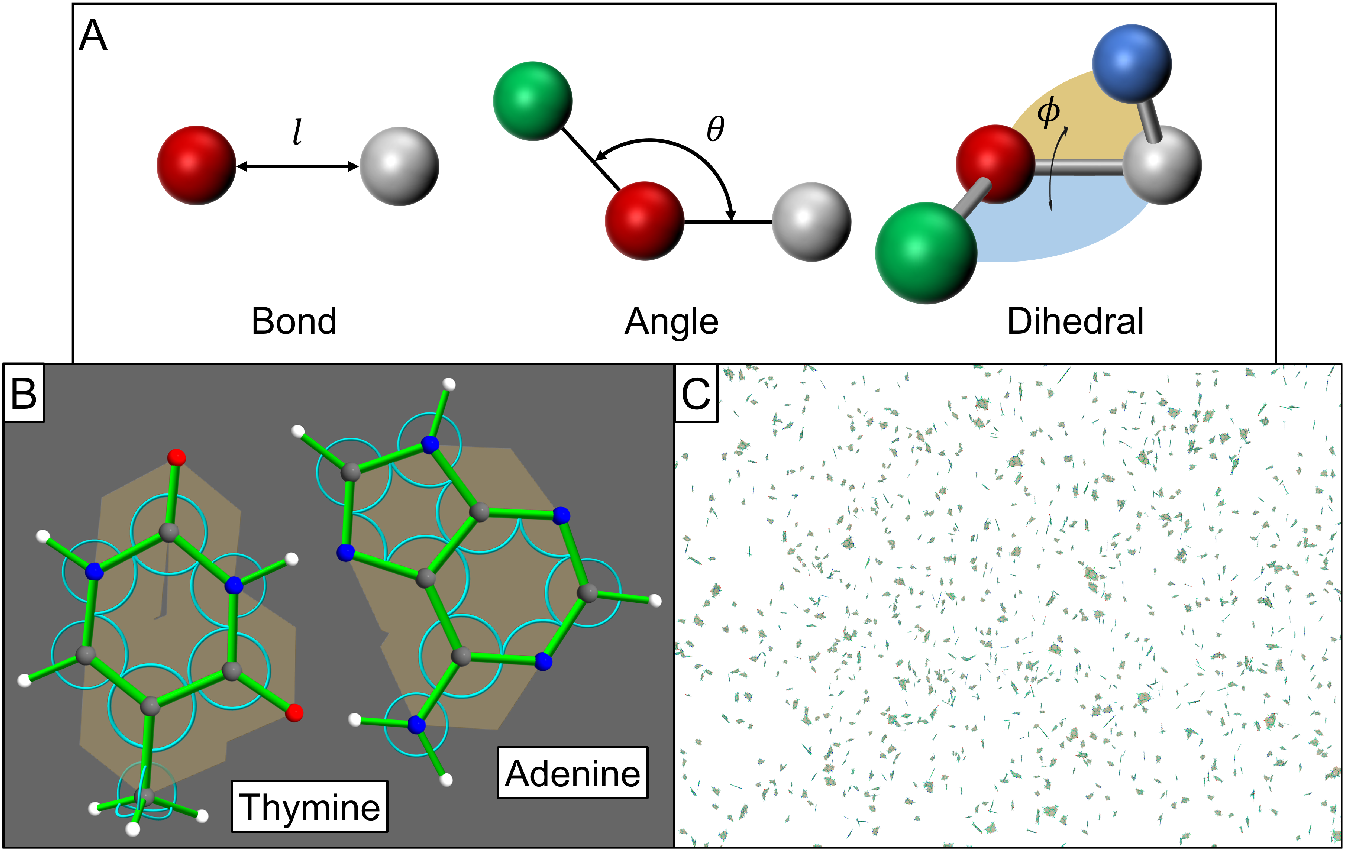
Molecular modeling and simulation with Tissue Forge. A: Classes of bonded interactions, where a measured property of the bond (length *l* for Bonds, angle *θ* for Angles, and planar angle *ϕ* for Dihedrals) is used as input to a potential function. B: Detailed view of thymine (left) and adenine (right) molecules constructed from Tissue Forge objects. Bonds shown as green cylinders, angles as blue arcs, and dihedrals as gold planes. C: Real-time simulation of a cloud of thymine and adenine molecules interacting via long-range potentials in a neutral medium.

Tissue Forge supports combining aspects of object-oriented programming with primitive Tissue Forge objects to define complex model objects for use in simulations. When modeling the dynamics of biomolecules, each particle can represent an atom, the atomic properties of which are defined through the Tissue Forge particle type. Definitions of particular biomolecules, such as nucleobases like thymine and adenine (Figure 4B) can then be designed using generic Python (or other supported language) classes that construct an instance of a biomolecule by assembling Tissue Forge particles and bonded interactions according to experimental data. Tissue Forge facilitates the construction and deployment of software infrastructure to develop interactive simulations of biomolecular systems and processes (Figure 4C, see Supplementary Materials S3).

Particle-based methods are also useful for coarse-grained modeling of subcellular components, where the atoms of individual biomolecules, biomolecular complexes, or even organelles are omitted and instead represented by a single particle that incorporates the aggregate behavior of its constituents (*e*.*g*., subcellular-element models). Tissue Forge supports coarse-grained subcellular modeling at various resolutions from the molecular to cellular scales, where a particle can represent a whole molecule, complex, or portion of an organelle or cytoplasm, to which coarse-grained properties (*e*.*g*., net charge or phosphorylation state) and processes (*e*.*g*., pumping of a solute, metabolism of a small molecule) can be applied.

For example, a particle can represent a portion of a lipid bilayer, in which case a sheet of such particle with appropriate binding and periodic boundary conditions can represent a section of a cell membrane. The Tissue Forge simulation domain can describe representative local spatial dynamics of the cell interface with its surrounding environment. Tissue Forge supports particle-based convection, providing a straightforward way to simulate a coarse-grained model of active transport at the cell membrane. Tissue Forge provides additional transport laws to model active pumping of species into or out of particles. To model transport at the cell membrane, these transport laws support implementing coarse-grain models of membrane-bound complexes like ion channels, which create discontinuities in concentrations of target species across the cell membrane (Figure 5, see Supplementary Materials S4).

**Figure 5:**
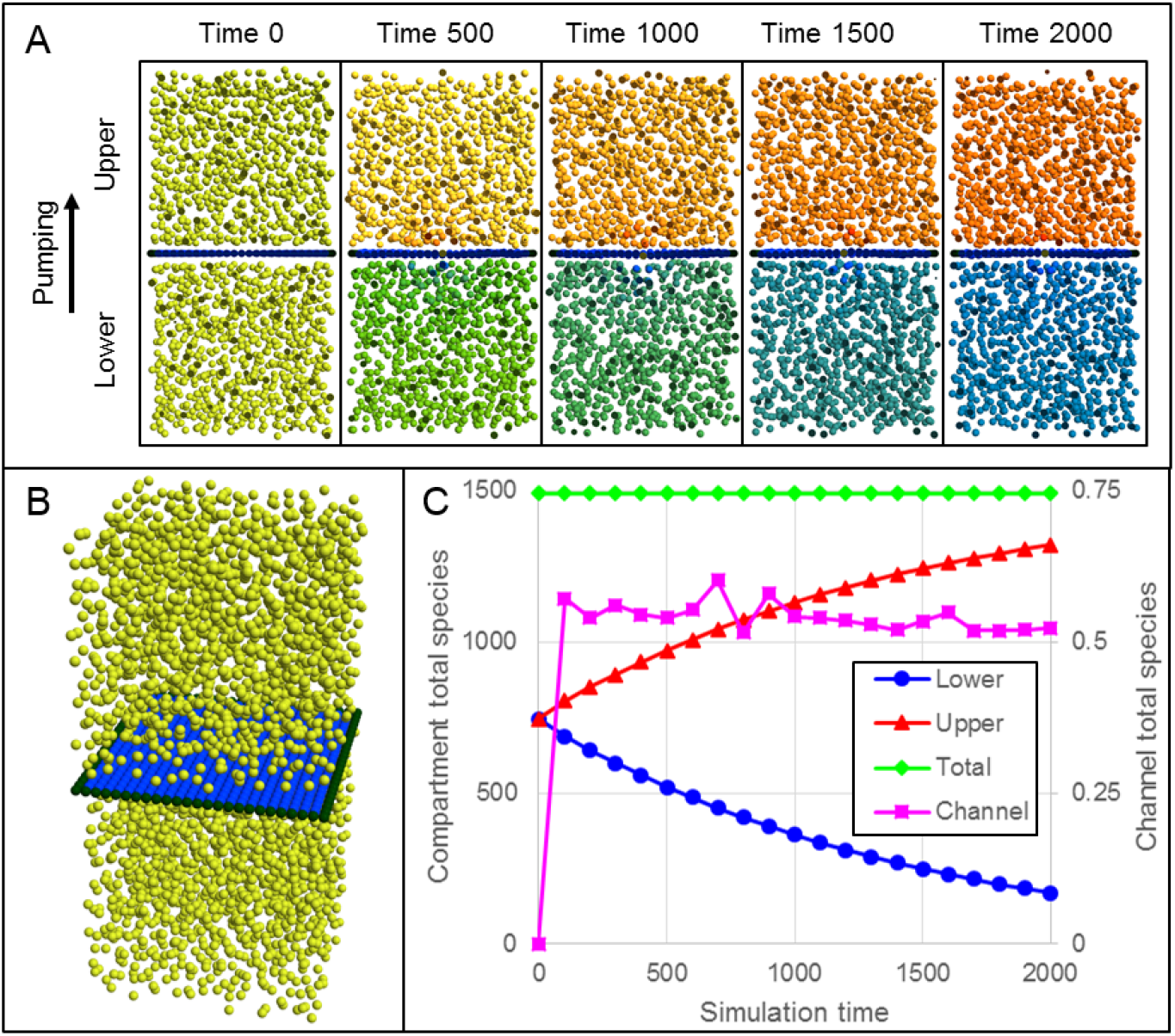
Active pumping of a diffusive species across a deformable membrane separating two fluid-filled compartments. A: Cut-plane views during simulation of two fluid-filled compartments separated by a deformable membrane, where each fluid is uniformly initialized with an initial concentration of a species. Particle color indicates species concentration with red as high, yellow and green as intermediate, and blue as low concentration. The membrane contains a particle that actively pumps the species from the lower to the upper compartment. B: Three-dimensional view of initial simulation state. C: Measurements of total species amounts in the lower (blue, circles), upper (red, triangles) and both (green, diamonds) compartments (left-hand vertical axis), and in the channel (magenta, squares, right-hand axis), during simulation.

At the coarsest scale of target applications, Tissue Forge provides support for particle-based modeling of multicellular dynamics. Tissue Forge provides a number of modeling features to support multicellular modeling at resolutions at or near the multicellular scale, where a particle can represent an individual cell, or a part of a cell. Overdamped dynamics describe the highly viscous, fluidlike collective motion of particle-based model cells, where short-range, implicit interactions can represent volume exclusion and contact-mediated intercellular interactions (*e*.*g*., adhesion), long-range, implicit interactions can represent intercellular signaling via soluble signaling, and particle state vectors can describe the intracellular state.

For example, particle-based model descriptions have been previously used to describe cells as a set of particles (*e*.*g*., a Tissue Forge cluster, Figure 1B) when modeling the process of spheroid fusion in tissue bioprinting, where cohesive cell shape is maintained by Lennard-Jones and harmonic potentials between particles of the same cell, and intercellular adhesion occurs by a Lennard-Jones potential between particles of different cells [12]. In a simpler model, representing each cell as a single particle and intercellular interactions with a single Morse potential can also produce emergent fusion of spheroids like those used in bioprinting of mineralized bone (*i*.*e*., about 12.5k cells per spheroid, Figure 6) [14]. When coupled with modeling diffusive transport and uptake like the scenario demonstrated in Figure 5, a Tissue Forge-based framework for the simulation of nutrient availability during spheroid-dependent biofabrication could support detailed modeling of spheroid viability in large tissue constructs [15].

**Figure 6:**
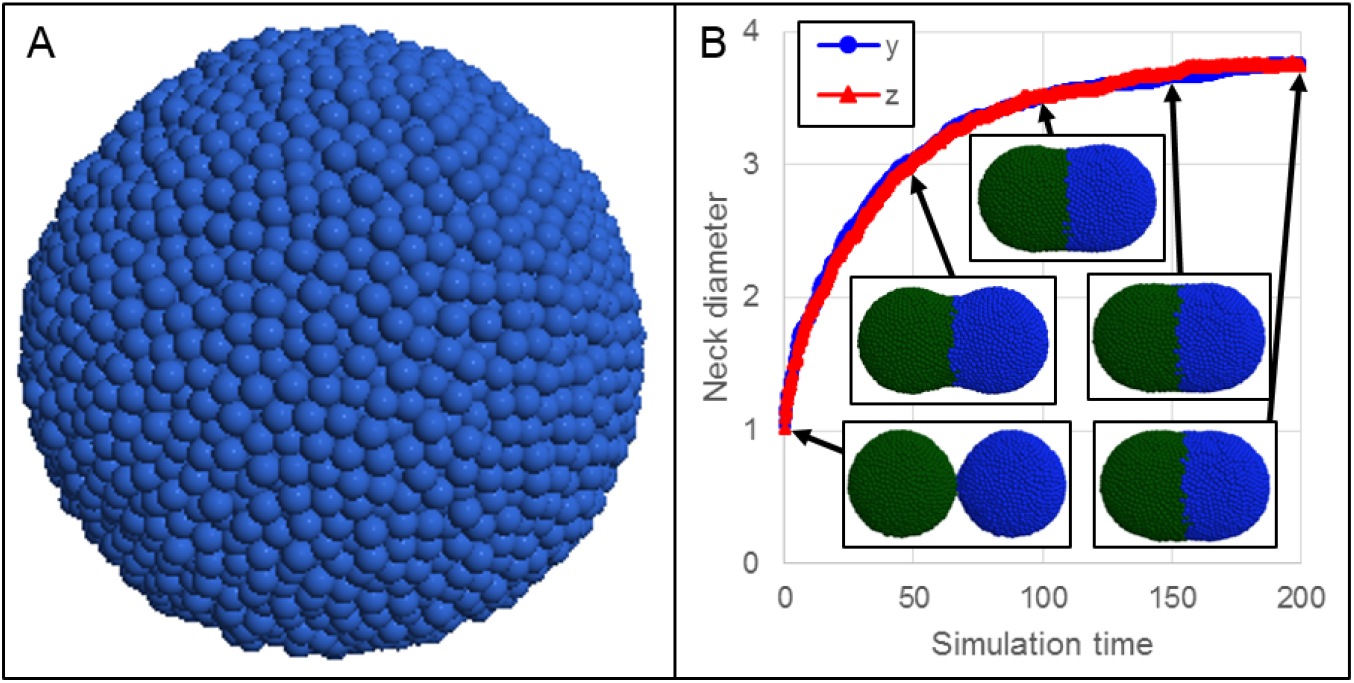
Simulating fusion of multicellular, homotypic spheroids. A: Spheroids of 12.5k cells each were individually pre-assembled, as in typical bioprinting practice. B: Two spheroids (green and blue) placed in close proximity fuse over time, as measured by the neck diameter along the *y* (blue circles) and *z* (red triangles) directions, which grows over time. The neck diameter along a direction is measured as the largest distance along the direction between any two particles at the mid-plane. Insets show the simulation at times 1, 50, 100, 150 and 200.

## 4 Discussion

The Tissue Forge modeling and simulation framework allows users to interactively create, simulate and explore models at biologically relevant length scales. Accessible interactive simulation is key to increasing scientific productivity in biomodeling, just as simulation environments are fundamental to other fields of modern engineering. Tissue Forge supports both interactive runs with real-time visualization for model development, and headless execution for data generation and integrated applications. In addition, Tissue Forge supports user-specified model features (*e*.*g*., custom particle types, forces and potentials) and scheduled and keyboard-driven simulation events, with intuitive user interfaces, in multiple programming languages and frameworks, supporting beginnerto expert-level programmers and beginnerto expert-level biomodelers.

Tissue Forge is open-source and freely available under the LGPL v3.0 license (https://github.com/tissue-forge/tissue-forge). Pre-built binaries are available in C, C++ and Python on 64-bit Windows, MacOS and Linux systems via conda (https://anaconda.org/tissue-forge/tissue-forge). Online documentation provides information on project philosophy, installation, walk-throughs, examples (in Jupyter Notebooks, https://github.com/tissue-forge/tissue-forge/tree/main/examples/py/notebooks) and API documentation for all supported languages. It has automated build updates to maintain synchronization between software versions and documented features (https://tissue-forge-documentation.readthedocs.io), including details on features not described in this paper (*e*.*g*., species transport, boundary conditions). Tissue Forge’s transparent development cycle, with automated continuous integration and continuous delivery, rapidly and reliably delivers the latest features to users (https://dev.azure.com/Tissue-Forge/tissue-forge). Instructions for installing Tissue Forge are available in the Supplementary Materials S1.

Tissue Forge applies the abstraction of a particle to support modeling applications over a wide range of scales, ranging from sub-nanometer to hundreds of micrometers and beyond. It supports future development and integration of advanced numerical and computational methods for incorporating and/or generating biological information with increasingly greater detail. Tissue Forge provides a designated space for development of application-specific models and methods by both the development team and user community, and so is free to grow and evolve into other computational domains with significant relevance and impact to a number of scientific communities. To this end, we are preparing a followup manuscript that demonstrates advanced modeling and simulation features, detailed model construction in specific applications, and relevant features that are currently under development. Tissue Forge features under development include improvements to core Tissue Forge simulation capability (*e*.*g*., multi-GPU support and libRoadRunner [16] integration for network dynamics modeling), additional modeling features (*e*.*g*., new built-in potentials and forces, support for improper angles in MD modeling), enhanced user experience (*e*.*g*., a graphical event interface), and additional modeling methodologies and solvers (*e*.*g*., vertex and subcellular element models).

## 5 Conclusion

Tissue Forge supports biological, chemical and physics research by providing an interactive modeling and simulation environment for particle-based model development, execution and sharing, including integration with applications in multiple programming languages. The Tissue Forge Python API supports interactive modeling as a standalone application or in a Jupyter Notebook, while the Tissue Forge C and C++ APIs support development of compiled and integrated applications for advanced and compute-intensive projects. Tissue Forge supports modeling applications over a broad range of scales, from the molecular to the multicellular and beyond, and adopts a robust architecture to grow according to the needs of target scientific communities.

## 6 Acknowledgments

Funding for Tissue Forge is provided by NIBIB U24 EB028887 (HMS, JAG, TJS, JPS). TJS and JAG acknowledge funding from grants NSF 2120200, NSF 2000281, NSF 1720625, NIH R01 GM122424. JPS acknowledges additional funding from the EPA STAR RD840027 and NSF 2054061. This research was supported in part by Lilly Endowment, Inc., through its support for the Indiana University Pervasive Technology Institute.

## 7 Author Contributions

**Conceptualization:** TJS, JPS, HMS, JAG

**Data Curation:** TJS

**Formal Analysis:** TJS

**Funding Acquisition:** TJS, JPS, HMS, JAG

**Investigation:** TJS

**Methodology:** TJS, JPS, HMS, JAG

**Project Administration:** TJS, HMS, JAG

**Resources:** TJS, JPS, HMS, JAG **Software:** TJS

**Supervision:** TJS, HMS, JAG

**Validation:** TJS **Visualization:** TJS

**Writing – Original Draft Preparation:** TJS, JPS, HMS, JAG

**Writing – Review Editing:** TJS, JPS, HMS, JAG

## 8 Supplementary Materials

**S1 Installing Tissue Forge**. Instructions for installing pre-built Tissue Forge binaries.

**S2 oscillator.ipynb**. Jupyter Notebook that simulates a simple oscillator with two particles.

**S3 dna.py**. Python script that constructs adenine and thymine nucleobases on the basis of individual atoms using Tissue Forge particles.

**S4 membrane.ipynb**. Jupyter Notebook that simulates a neighborhood at a deformable membrane separating two fluids and active transport between them.

